# Low-intensity vibration does not induce changes in microtubule dynamics in vitro

**DOI:** 10.1101/2024.03.20.585904

**Authors:** Chase A. Crandall, Nina N. Nikitina, Anamaria G. Zavala, Sean M. Howard, Gunes Uzer

## Abstract

Microtubules (MTs) are cytoskeletal filaments responsible for many vital cellular processes including intracellular organelle organization and enabling the movement of intracellular components. While MTs were shown to respond to low frequency and large mechanical signals like substrate strain, how MTs may respond to a high frequency mechanical signal like low-intensity vibrations (LIV) is unknown. Here we quantified the polymerization dynamics of MTs under an acute 1-day LIV protocol applied at 90 Hz and 0.7 xg, a signal we have shown to be effective for altering F-actin dynamics and nuclear stiffness. LIV treatments were compared against Taxol, a potent regulator of MT acetylation. Using mouse mesenchymal stem cells (MSCs) *in vitro*, we quantified tubulin polymerization via centrifugal fractionation and western blots as well as alpha-tubulin acetylation via immunostaining. Finally, MT growth dynamics were quantified using machine learning-assisted analysis of live cell fluorescence microscopy of MT plus end binding protein EB1. Our results were not able to detect differences between LIV and control groups while Taxol treatment was effective in all measured outcomes. Our findings indicate that LIV applied at 90 Hz and 0.7 xg does not affect MT dynamics in MSCs, suggesting a higher mechanical threshold of MTs when compared to F-actin cytoskeleton.

## Introduction

In the field of mechanobiology, the relationship between mechanical forces and remodeling of cell cytoskeleton is central. Dynamic remodeling of the cytoskeleton informs MSCs, which play a pivotal role in tissue regeneration and repair^1,2^, how to behave and evolve based on the physical signals they receive. These cells interpret mechanical cues from the surroundings, guiding their differentiation and function–a process impacted by cytoskeletal dynamics, including those of both actin filaments and MTs^3^.

At the cellular level, LIV promotes focal adhesion activation^2^ and βcatenin signaling^4^, YAP nuclear entry^5^, cell proliferation^6^, and accelerates osteogenesis of MSCs^7^. The effects of LIV on cell function are, at least in part, modulated by changes in cell structure. LIV increases RhoA-induced contractility, resulting in stiffer F-actin struts^8^ and increases nuclear stiffness^9^. While the role of LIV on MT dynamics is currently unknown, the interplay between MT and the actin cytoskeletons is integral to cell function, including sustaining cell shape, enabling cell movement division, and facilitating intracellular transport^10,11^. Thus, the intricate relationship between actin filaments and MTs—where mechanical stimuli affecting one component can lead to compensatory or parallel changes in the other^12^—raises a critical question: do MTs exhibit similar responsiveness to LIV as observed with actin?

MTs display dynamic instability, a behavior characterized by alternating periods of growth and shrinkage. MT growth is partially coordinated by +TIPs—proteins that are primarily recruited by end-binding proteins (EBs) to the growing end of MTs, thus enabling precise measurement of their dynamics^13^. The dynamic nature of MTs disposes them to their ability to respond to a range of mechanical stimuli, such as stretching, compression, and modulations of the extracellular matrix^14^. However, the specific response of MTs to LIV has yet to be comprehensively explored. Therefore, to further characterize a component of the cellular mechanotransduction pathway that remains poorly understood, this study investigated the influence of LIV on MT dynamics *in vitro*. To that end, we applied LIV to mouse MSCs and quantified the effects on MTs using western blot analysis of tubulin polymerization, immunocytochemical analysis of alpha-tubulin acetylation, and live cell fluorescence microscopy of MT plus end-binding protein, EB1, with associated computational analysis and machine learning to quantify MT growth dynamics.

## Materials and Methods

### Cell Culture

Bone marrow derived MSCs from 8–10 wk male C57BL/6 mice were isolated as described from multiple mouse donors, and MSCs pooled, providing a heterogenous MSCs cell line^15^. Briefly, tibial and femoral marrow were collected in RPMI-1640, 9% FBS, 9% HS, 100 μg/ml pen/strep, and 12 μM L-glutamine. After 24 hours, non-adherent cells were removed by washing with phosphate-buffered saline (PBS), and adherent cells were cultured for 4 weeks. Passage 1 cells were collected after incubation with 0.25% trypsin/1 mM EDTA × 2 minutes and re-plated in a single 175-cm^2^ flask. After 1–2 weeks, passage 2 cells were re-plated at 50 cells/cm2 in expansion medium (Iscove’s modified Dulbecco’s Medium (IMDM), 9% FBS, 9% HS, antibiotics, L-glutamine). MSCs were re-plated every 1–2 weeks for two consecutive passages up to passage 5 and tested for osteogenic and adipogenic potential, and subsequently frozen.

### Experimental Design

MSCs derived from mouse bone marrow were plated and cultured in IMDM with 10% fetal calf serum (FCS) and 1% Penicillin-Streptomycin (P/S). Low-intensity vibration (LIV) was applied in 6 rounds. In each round, plates were placed on a machine at room temperature (RT) that vibrated them at 90 Hz, 0.7 xg for 20 minutes. Control plates were not vibrated but were kept at RT while the other plates vibrated. Plates were incubated for 60 minutes at 37 °C between each round. Samples were taken 3 hours following the final round of LIV.

### Centrifugal Separation of Polymerized and Unpolymerized MT

MSCs were plated at 100K/well on 6-well plates (see Figure 1a). Cells were incubated at 37 °C for ∼30 hours before being starved using IMDM with 2% FCS, 1% P/S, 5 ug/mL insulin, 0.1 µM dexamethasone, and 50 µM indomethacin. Cells remained in this media for ∼12 hours before the start of vibration. LIV was applied as described above, and samples were collected using the “Microtubules/Tubulin In Vivo Assay Kit” (Cytoskeleton, Cat. #BK038) as per the manufacturer’s instructions. Briefly, lysates were taken down into a MT-stabilizing buffer. The lysates were homogenized and centrifuged at 1000 xg for 5 minutes at 37 °C to obtain the low-speed pellet (LSP) containing the large MTs. 100 µL of the supernatant from each sample was centrifuged at 100,000 xg for 60 minutes at 37 °C to obtain the high-speed pellet (HSP) containing the small MTs and the high-speed supernatant (HSS) containing free tubulin. The LSP, HSP, and HSS were analyzed for tubulin content by western blot as per the manufacturer’s instructions to determine the fraction of the cell’s tubulin present in the form of large MT, small MT, and free tubulin, respectively (see Equation 1). A positive control was performed using Taxol. Cells were treated with Taxol (3 µM)^16^ in DMSO and incubated for 3 hours before samples were collected as described above. They were compared with corresponding controls treated solely with DMSO.

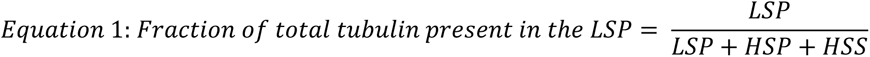

**Figure 1:**
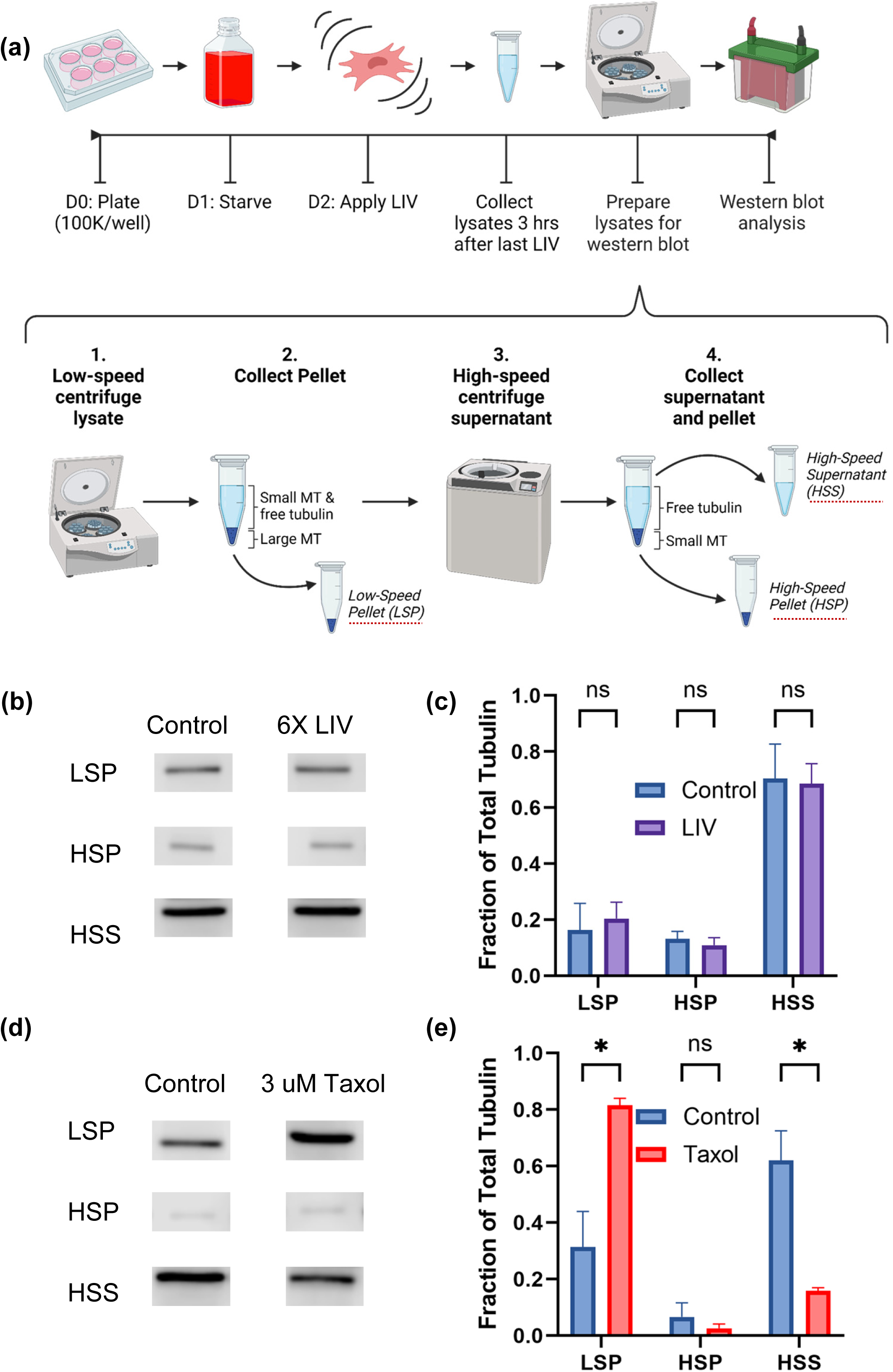
**(a)** Diagram of experimental workflow for determining tubulin polymerization. **(b)** Western blot analysis depicting the total tubulin content in the three fractions—Low-Speed Pellet (*LSP*), High-Speed Pellet (*HSP*), and High-Speed Supernatant (*HSS*)— for both LIV (Low-Intensity Vibration) treated and Control cells. **(c)** Bar graph representation of the tubulin content determined by western blot analysis in the three fractions for LIV and Control cells. The mean percentage of tubulin content in each fraction for Control cells was found to be: LSP 0.3138, HSP 0.06605, HSS 0.6201. For LIV treated cells, no significant difference in tubulin content was observed in comparison to Control cells (p>0.05), suggesting that LIV does not significantly alter tubulin polymerization. **(d)** Western blots showing total tubulin content in each of the three fractions–LSP, HSP, and HSS–for LIV and Control cells. (e) Bar graph showing tubulin content determined by western blot analysis in each of the three fractions for LIV and Control cells. There was no significant difference between LIV and Control cells. Full–uncut plots are presented in Figures S1-S6. **(e)** Positive control analysis via bar graph, comparing the mean percentage of tubulin in the three fractions for Taxol and Control cells. Control cells exhibited the following mean percentages: LSP 0.1638, HSP 0.1317, HSS 0.7045. Taxol-treated cells showed: LSP 0.2047, HSP 0.11, HSS 0.6853. A paired t-test revealed a significant difference in the LSP and HSS fractions between the two groups (p=0.023 and p=0.0198 respectively), suggesting that Taxol effectively increases polymerized tubulin in the LSP while reducing free tubulin in the HSS.

### ICC Analysis of Alpha-Tubulin Acetylation

MSCs were plated at 7.5K per well in IBIDI 35 mm imaging plates, then incubated, starved, and vibrated as described above (see Figure 2a). 3 hours after the final round of LIV, cells were fixed in warm formaldehyde (4%, 37 °C) for 10 minutes at RT. Plates were washed twice for 3 minutes in ice-cold PBS before being permeabilized in Triton (0.3%) for 15 minutes at RT. Plates were successively incubated in Anti-Acetyl-alpha-Tubulin (Lys40) rabbit monoclonal antibody (clone: RM318, 1:1000) for 60 minutes at 37 °C, AlexaFluor594 Goat Anti-Rabbit antibody (1:300) for 30 minutes at 37 °C, and a Hoechst 33342 stain (0.5 drops/mL) for 20 minutes at RT. Plates were then protected from photobleaching by addition of an antifade mounting medium with a coverslip. Plates were washed 3 times for 3 minutes with ice-cold PBS between incubations unless otherwise noted.

**Figure 2:**
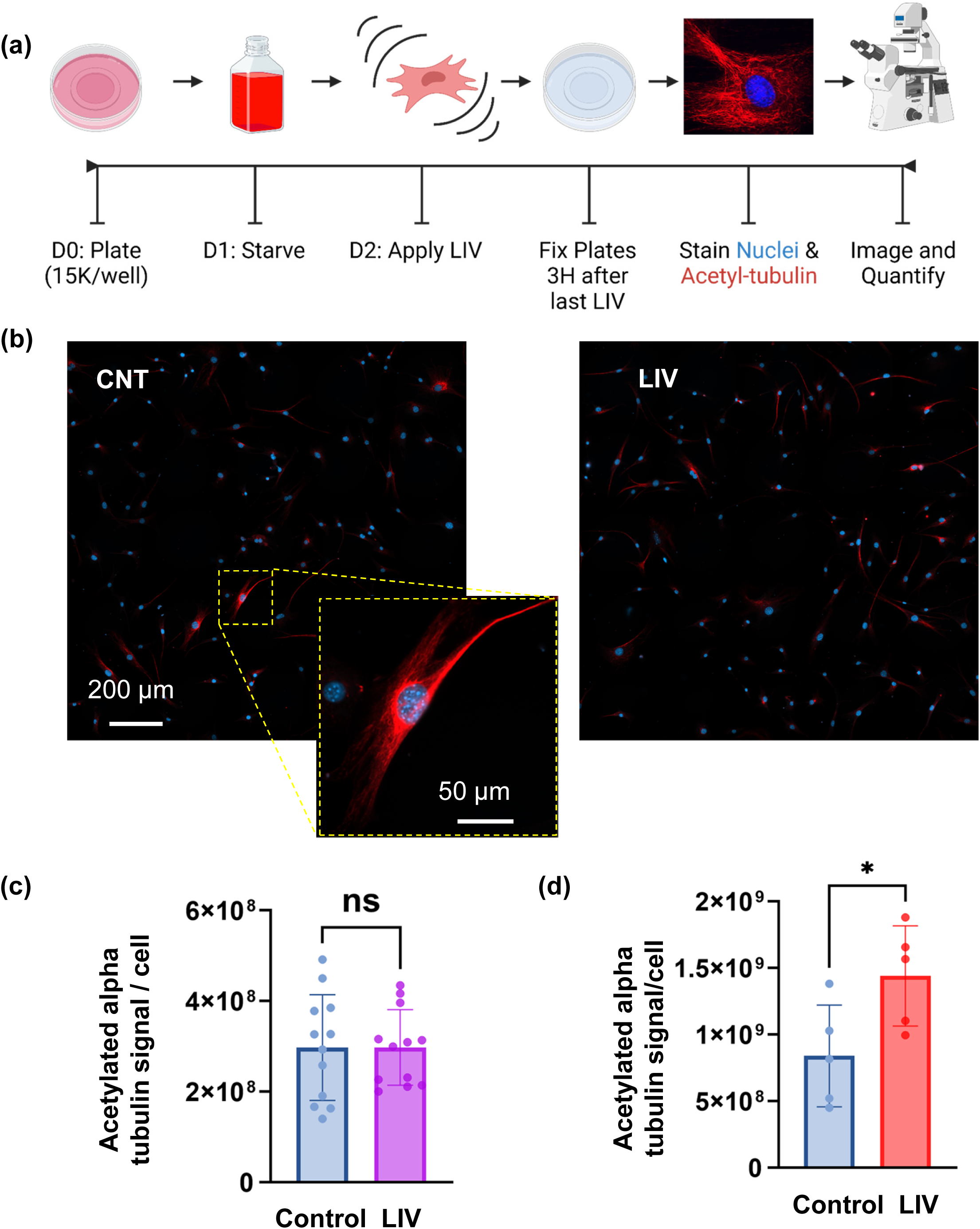
**(a)** Diagram of experimental workflow for immunocytochemical analysis of alpha-tubulin acetylation. **(b)** Representative fluorescence microscopy images display the cellular nuclei in blue and acetylated (K40) alpha-tubulin. Images were captured using a 40x oil immersion lens. Multiple images were stitched together to create a larger composite image (5×5 tiles), allowing for the visualization of a greater number of cells at this magnification level. **(c)** Acetylated Alpha-Tubulin Signal Quantification in LIV and Control Cells: Scatter plots present the quantification of the acetylated (K40) alpha-tubulin signal intensity per cell. Mean values and standard deviations are depicted for both LIV-treated and control cells. Statistical analysis indicated no significant difference in acetylated tubulin levels between the two groups (p>0.05), suggesting that LIV treatment does not markedly influence tubulin acetylation under these experimental conditions. **(d)** Acetylated Alpha-Tubulin Signal Quantification in Taxol and Control Cells: scatter plots show the acetylated (K40) alpha-tubulin signal per cell for Taxol-treated and control groups. Significant differences being noted between the two groups (p<0.05).

The experiment was repeated with a Taxol control. The cells were plated at 10K in IBIDI 35 mm imaging plates, then incubated and starved as described above. In place of vibration, they were treated with Taxol (3 μM) in DMSO. They were compared with corresponding controls treated solely with DMSO. Samples were fixed and stained after a 3-hour incubation with Taxol or DMSO. Reagents used for immunofluorescence and their concentrations are listed in **Supplementary Table S1**.

Imaging was conducted using a Zeiss LSM900 Microscope with a 40x/1.4 Oil objective and a 5×5 tile setting. This configuration allowed for an extensive field of view per tiled image, with the 40x Oil objective facilitating optimal signal detection. Image analysis was performed using a custom-developed program that automatically extracted the total red fluorescence signal (indicative of acetylated K40 alpha-tubulin) and the number of nuclei on each image. This in-lab-created program, instrumental for the analysis, is available on GitHub at: https://github.com/mal-boisestate/Immunostained_Image_Analysis

The acetylated K40 alpha-tubulin signal was divided by the number of cells in each stitched image to calculate the average signal intensity per cell. The number of cells was determined based on the count of nuclei present in the images (Figure 2b, 2c)

### MT Growth Speed Determination using PlusTipTracker

MSCs were plated in a 6-well plate at 75K/well in IMDM with 10% FCS and 1% P/S (see Figure 3a). In order to visualize EB1 movement—and therefore MT growth—cells were transfected the following day with the plasmid, pEB1-2xEGFP^13^ using the Invitrogen Lipofectamine 3000 Transfection kit according to the manufacturer’s directions. 48 hours later, cells were selected with G418 (1 mg/mL). 48 hours later, cells were plated at 60K/well in IBIDI 35 mm plates in phenol red-free DMEM with 10% FCS and 1% P/S to allow for clearer imaging. 24 hours later, LIV was applied as described above. Cells were imaged during the 3 hours following the last vibration.

**Figure 3:**
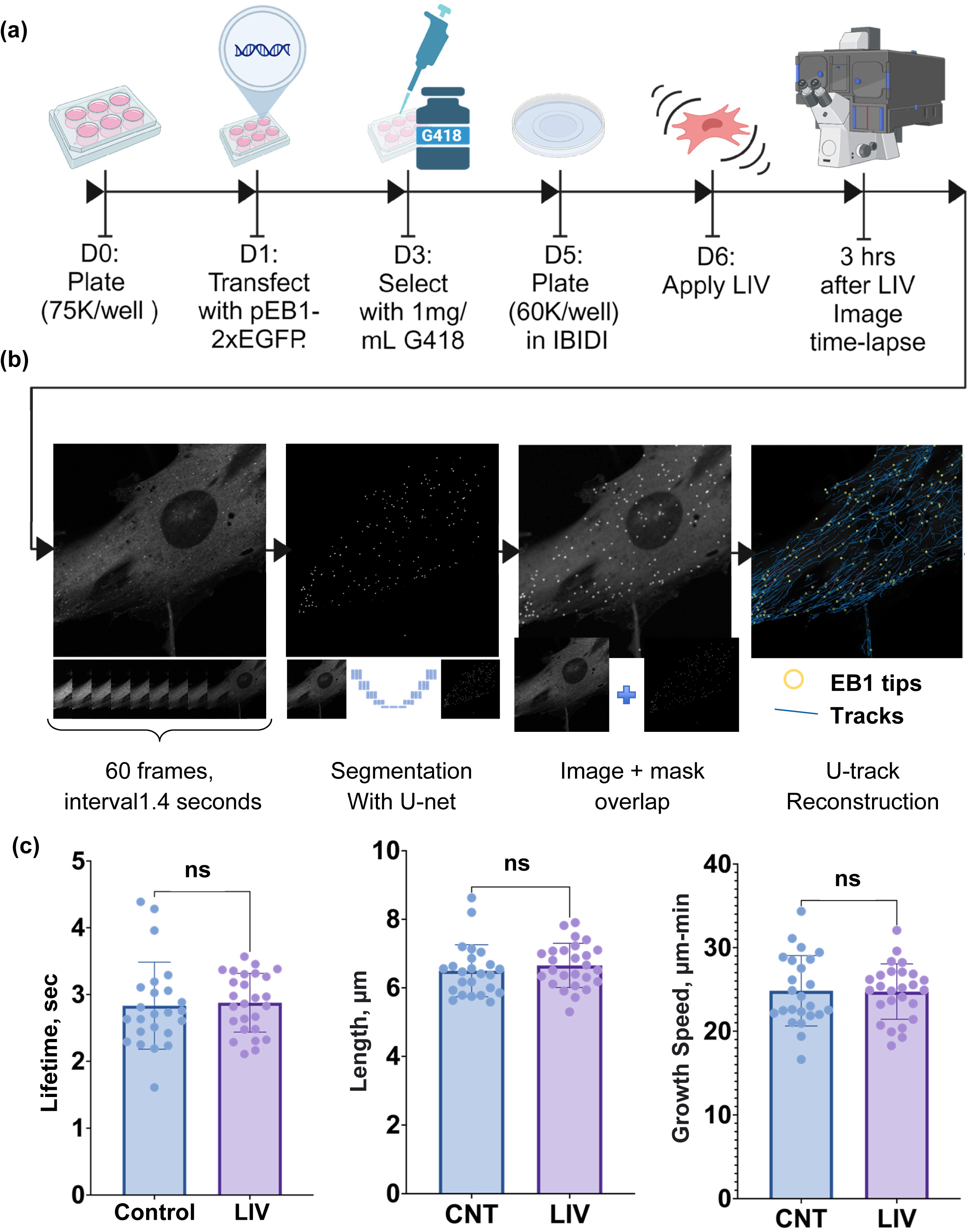
Microtubule Growth Speed Determination and Quantitative Analysis of Microtubule Dynamics Post-LIV Treatment. **(a)** Experimental Flow Illustration of MSC culture and transfection, selection with G418, plating in IBIDI plates, and LIV application. This followed by live cell imaging using a Zeiss LSM900 microscope, capturing 60 frames over an hour to create time-lapse movies for analysis with u-track 2.0. **(b)** PlusTipTracker (U-track) and Enhanced Imaging Analysis: processing workflow starting with time-lapse file ([Supplementary Movie 1](https://drive.google.com/file/d/1fz0Jyan0JkfJ6by3h_nWf1u8WGBB7rZq/view?usp=drive_link)), EB1 tip segmentation via a U-Net model for image enhancement ([Supplementary Movie 2](https://drive.google.com/file/d/1hLvggIqPdhAZlGPeyJweiDax6bLd9ZvR/view?usp=drive_link)) and, finally, EB1 track reconstruction through u-track MATLAB program ([Supplementary Movie 3](https://drive.google.com/file/d/1eCfhPSKkEhDsWkWjqgrL-7QR4NbPNTCn/view?usp=drive_link)). **(c)** Microtubule Dynamics Metrics: The scatter plots, with bars indicating mean and standard deviation, show microtubule lifetime, length, and growth speed. The dots on the scatter plots represent averaged data for specific cells, following initial individual calculations of growth speed, length, and lifetime. For the control group, the mean microtubule lifetime was 2.83 (Control) and 2.88 (LIV) seconds, the mean track length was 6.50 (Control) and 6.65 (LIV) micrometers, and the mean growth speed were 24.83 (Control) 27.74 (LIV) µm/minute. No significant differences were found in these parameters when comparing the LIV-treated cells to the control (p>0.05).

Utilizing a 40x oil objective on a Zeiss LSM900 microscope, live cell imaging was performed with uniform settings, followed by deconvolution. Each image formed part of a time-lapse sequence of 60 frames ([Supplementary Movie 1](https://drive.google.com/file/d/1fz0Jyan0JkfJ6by3h_nWf1u8WGBB7rZq/view?usp=drive_link)), creating an 84-second movie with intervals of 1.4 seconds between frames (see Figure 3b). The Plus-Tip Tracker software, now known as U-track 2.0, was sourced from the Danuser Lab GitHub page (https://github.com/DanuserLab/u-track)^17^. U-track 2.0 is an advanced version that includes the capabilities of the initially created PlusTipTracker program. For analysis, the software was applied to .czi files modified from the default settings: the difference of Gaussian filter parameters was set to a high of 3 pixels and a low of 5 pixels, watershed segmentation parameters adjusted to a minimum threshold of 6 standard deviations and a step threshold size of 4 standard deviations, tracking parameters specified with a maximum gap closure of 10 frames and a minimum track segment length of 3 frames.

Post-analysis, each reconstructed movie (47 for LIV and 53 for Control) underwent evaluation by researchers to track reconstruction quality on a scale of 1 to 10. Scores equal to or above 7 indicated precise reconstruction, while scores below were deemed poor. Initial results showed that 16 (34%) files from the LIV group and 29 (55%) from the control group scored above 7. The main challenge, leading to lower results, was distinguishing tips from noise, even with the advanced capabilities of PlusTipTracker.

A preprocessing step was employed to enhance tip detection, integrating a trained U-Net^18^ architecture-based convolutional neural network for segmenting EB1 tips on each image frame. Training and validation sets were manually created by labeling each EB1 tip dot. From 11 cell images, 30 frames were randomly assigned for training (24 frames) and validation (6 frames). The images were resized to 512×512 pixels for training. The learning parameters included a learning rate of 0.001, a batch size of 1, and a total of 200 epochs. A GPU was utilized to expedite the training process. The network was trained to minimize a loss function that quantifies pixel-to-pixel differences between predicted and target images, with the loss function adjusted to heighten the penalty for false negatives in actin fibers to 20.

Segmented tips from the U-Net model output were overlaid onto original images ([Supplementary Movie 2](https://drive.google.com/file/d/1hLvggIqPdhAZlGPeyJweiDax6bLd9ZvR/view?usp=drive_link)) and fed into U-track 2.0 as a set of TIFF files named according to U-track documentation for further EB1 tips track reconstruction ([Supplementary Movie 3](https://drive.google.com/file/d/1eCfhPSKkEhDsWkWjqgrL-7QR4NbPNTCn/view?usp=drive_link)).

Watershed segmentation parameters were fine-tuned to a minimum threshold of 7 standard deviations and a threshold size of 5 standard deviations, accommodating the slightly larger segmented tips. This adjustment changed the number of images with high-quality tracks to 26 (55%) for the LIV group and 24 (45%) for the control group, indicating improved detection accuracy and track reconstruction.

A series of supplementary movies compare track reconstruction across the time-lapse sequence based on raw and U-net EB1 comet segmented images: [Supplementary Movie 4](https://drive.google.com/file/d/1NK16VC3mLIOC8aBnNqbrz1BM1JBB-5Ju/view?usp=drive_link) juxtaposes raw and enhanced time-lapse, and [Supplementary Movie 5](https://drive.google.com/file/d/1NzXTUaoNHpHFWkomnuPFggOkXN9djHh6/view?usp=drive_link) demonstrates the improvement in the detection of EB1 comets by U-track 2.0. Finally, [Supplementary Movie 6](https://drive.google.com/file/d/1Q4aNsg1oEYPmJmVlOZEonZzrW5HV8b21/view?usp=drive_link) illustrates the improvement in track reconstruction performance with U-Net, where an image previously excluded due to the reconstruction’s low quality is recovered and analyzed successfully.

U-track software quantified MTs’ lifetime, length, and growth speed in each image. To decrease the false positive track number, we applied a minimum track length threshold of 2 µm, filtering out shorter tracks likely caused by noise. Since each image featured a single cell, we averaged the track metrics per image, yielding cellular-level data. Using the Mann-Whitney statistical test, we then compared these metrics for the LIV and Control groups.

## Results

### Taxol treatment but not LIV increases tubulin polymerization and alpha-tubulin acetylation

We first assessed the effect of LIV on tubulin polymerization (Figure 1). Through western blot analysis, we compared the tubulin content in three cellular fractions: the Low-Speed Pellet (LSP), High-Speed Pellet (HSP), and High-Speed Supernatant (HSS) (Figure 1b) that we got through Centrifugal Separation of Polymerized and Unpolymerized MT described in the method section (Figure 1a). Our results demonstrated that LIV treatment did not significantly alter the distribution of tubulin among these fractions when compared to control cells (Figure 1b, 1c); there is no significant difference observed in tubulin content between LIV-treated and control cells for all three fractions(p>0.05) (see Figures S1, S2, S3 for full blots). As a positive control, we compared the tubulin content in Taxol-treated and control cells (Figure 1d). Taxol, a known MT stabilizer, significantly increased polymerized tubulin in the LSP while reducing free tubulin in the HSS (Figure 1e), validating our experimental approach (see Figures S4, S5, S6 for full blots).

Investigating alpha-tubulin acetylation, we performed immunocytochemical assays and analyzed the resultant images to assess acetylation levels (Figure 2a). Fluorescent microscopy was used to get images and then stitch them together (Figure 2b), thereby providing an extensive field of view. Our analysis encompassed 12 tiled images each for the LIV and Control groups, containing a total of 1512 and 1665 cells, respectively. The acetylation signal was divided by the number of cells in each image. Statistical evaluation revealed no significant alteration in acetylation (K40) of alpha-tubulin in the LIV-treated group compared to the control (p>0.05) (Figure 2c). This result suggests that LIV exposure does not significantly influence the acetylation state of alpha-tubulin.

In contrast, the Taxol-treated group showed a noticeable increase in alpha-tubulin acetylation levels when compared to the control, with the analysis of 5 tiled images per group—comprising 739 cells in the control group and 590 in the Taxol-treated group—revealing significant differences (p=0.0366) (Figure 2d). This disparity serves as a positive control, affirming our experimental setup’s sensitivity and discriminative capability to detect acetylation changes induced by Taxol.

### LIV does not alter MT dynamics

Finally, from fluorescent live-cell time-lapse movies of EB1 on growing ends of MTs, we extrapolated and quantified MT dynamics metrics in post-LIV treatment (Figure 3) using computational analysis with U-track 2.0 enhanced by deep learning preprocessing step. Metrics such as MT lifetime, length, and growth speed showed no significant differences between LIV-treated cells and controls (p>0.05). Specifically, the mean MT lifetime was 2.83 seconds for the control group and 2.88 seconds for the LIV group, the mean track length was 6.50 micrometers (Control) versus 6.65 micrometers (LIV), and the mean growth speed was calculated at 24.83 µm/minute (Control) and 27.74 µm/minute (LIV).

## Discussion

In this study, we have analyzed MTs via a multitude of methods - employing protein quantification assays, immunocytochemical techniques, live cell fluorescence microscopy, and image analysis with machine learning. These methods showed no significant differences in tubulin polymerization, alpha-tubulin (Lys40) acetylation, MT growth speed, length, or lifetime between the LIV and control groups. This outcome, supported by agreement between all analysis methods, shows a resilience of MT dynamics to LIV–in contrast to the actin cytoskeleton’s responsiveness. This suggests a fundamental divergence in how MTs and actin filaments respond to mechanical cues. Research, including strain experiments by Kubitschke *et al*. (2017)^19^, highlights this differential responsiveness, indicating that MT and actin filament systems may be tuned to distinct magnitudes of mechanical signals. Further, in light of a study by Putnam, *et al*. (2001)^14^ that found MTs react oppositely to compressive and tensile forces and the fact that MTs align along the axis of maximal stress (Hamant 2008)^20^, the application of LIV might be read by cells as alternating equal and opposite–and therefore cancellatory–tensile and compressive forces.

The limitations of our experimental methodologies could also help explain the lack of MT response to LV. The fact that no significant differences in MT mass ratios were found between LIV and control could be attributed to the precision of the assay. One compounding variable was temperature. According to the centrifugal separation kit’s manufacturer, a 1°C drop in temperature in any step of the protocol can result in a 5% decreased MT mass. Even though LIV and control samples were collected in parallel, the day-to-day variability in the temperature of the lab could have contributed to the error, obscuring potential differences between treatments. However, our Taxol controls seem to be not affected by this potential variability. Our other assays should have significantly better precision. However, the possibility remains that the conditions under which LIV was applied did not meet the threshold necessary to prompt a detectable response in MT dynamics.

Considering the critical roles of MTs in cellular architecture and signaling, understanding the mechanistic basis of their stability or adaptability under mechanical stress remains a compelling area for future research. Future studies could examine the effects of increasing LIV intensity, frequency, and duration. An in-depth exploration of alterations in MT structural organization could also further characterize MT mechanoresponse: identifying specific cellular regions more prone to changes and determining whether the MT network becomes denser and more interconnected. Such studies deepen our understanding of the cytoskeletal interplay, potentially revealing new aspects of cellular mechanobiology.

While our study did not find evidence of LIV-induced alterations in MT dynamics, the lack of response deepens our understanding of cytoskeleton mechanoresponse and helps to direct further research in this field.

## Acknowledgements

pEB1-2xEGFP was a gift from Torsten Wittmann (Addgene plasmid # 37827 ; http://n2t.net/addgene:37827 ; RRID:Addgene_37827). This study was supported in part by NIH AG059923, NSF 1929188 and NSF 2025505. We acknowledge support from COBRE, INBRE, the Institutional Development Awards (IDeA) from the NIGMS Grants #P20GM103408, P20GM109095, and IC06RR020533. We also acknowledge support from the Biomolecular Research Center with funding from the NSF Grants #0619793 and #0923535; the M.J. Murdock Charitable Trust; Lori and Duane Stueckle, and the Idaho State Board of Education.

Figures 1a, 2a, and 3a were created with Biorender.

## Author contributions

Chase A. Crandall: concept/design, data analysis/interpretation, manuscript writing Nina N. Nikitina: concept/design, data analysis/interpretation, manuscript writing Anamaria G. Zavala: troubleshooting, guidance/supervision, final approval of manuscript Sean M. Howard: troubleshooting, guidance/supervision, final approval of manuscript Gunes Uzer: concept/design, data analysis/interpretation, financial support, final approval of manuscript

## Data availability

The datasets generated during and/or analyzed during the current study are available from the corresponding author upon reasonable request.

---

**Supplementary Table S1:** Table of reagents used for immunofluorescence and their concentrations.

| Reagent: | Concentration (in PBS): |
| --- | --- |
| Formaldehyde | 4% |
| Triton | 0.3% |
| Goat Serum | 5% |
| Hoechst 33342 stain | Concentration not disclosed by manufacturer; diluted 0.5 drops/mL |
| Anti-Acetyl-alpha-Tubulin (Lys40) rabbit monoclonal antibody (clone: RM318) | Concentration not disclosed by manufacturer; diluted 1:1000 |
| AlexaFluor594 Goat Anti-Rabbit antibody | Diluted 1:300 to a final concentration of 6.7 µg/mL |

## Supplementary Figures

**Figure S1:**
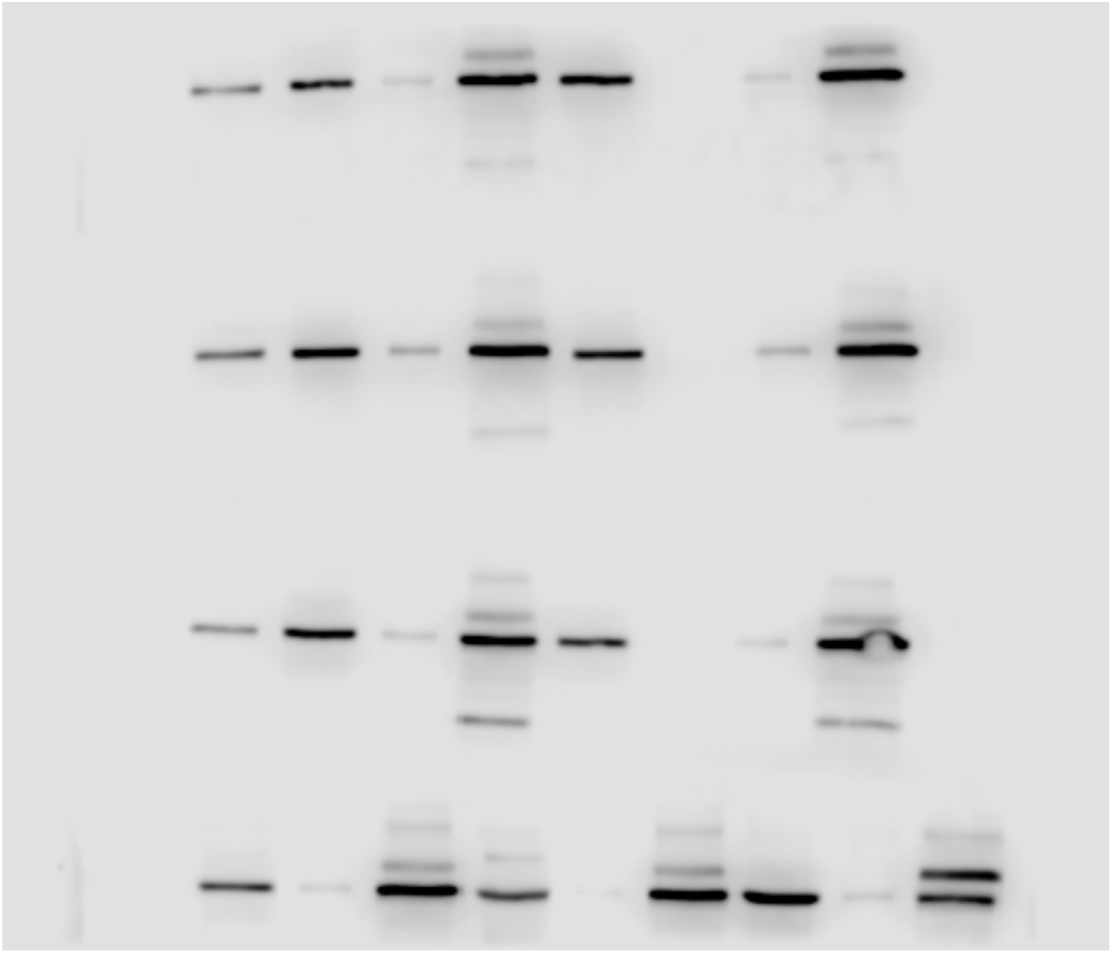
Taxol vs Control - Replicate 1. Western blot analysis of tubulin content present in the LSP, HSP, and HSS for Taxol-treated and control samples. The leftmost band of each row (except the 4th from the top) represents a 50 ng tubulin standard. Apart from that band, control samples are found in the left half of the 3rd row from the top, from left to right LSP, HSP, HSS. Taxol-treated samples are the rightmost 3 bands of the 4th row from the top, from left to right: LSP, HSP, HSS.

**Figure S2:**
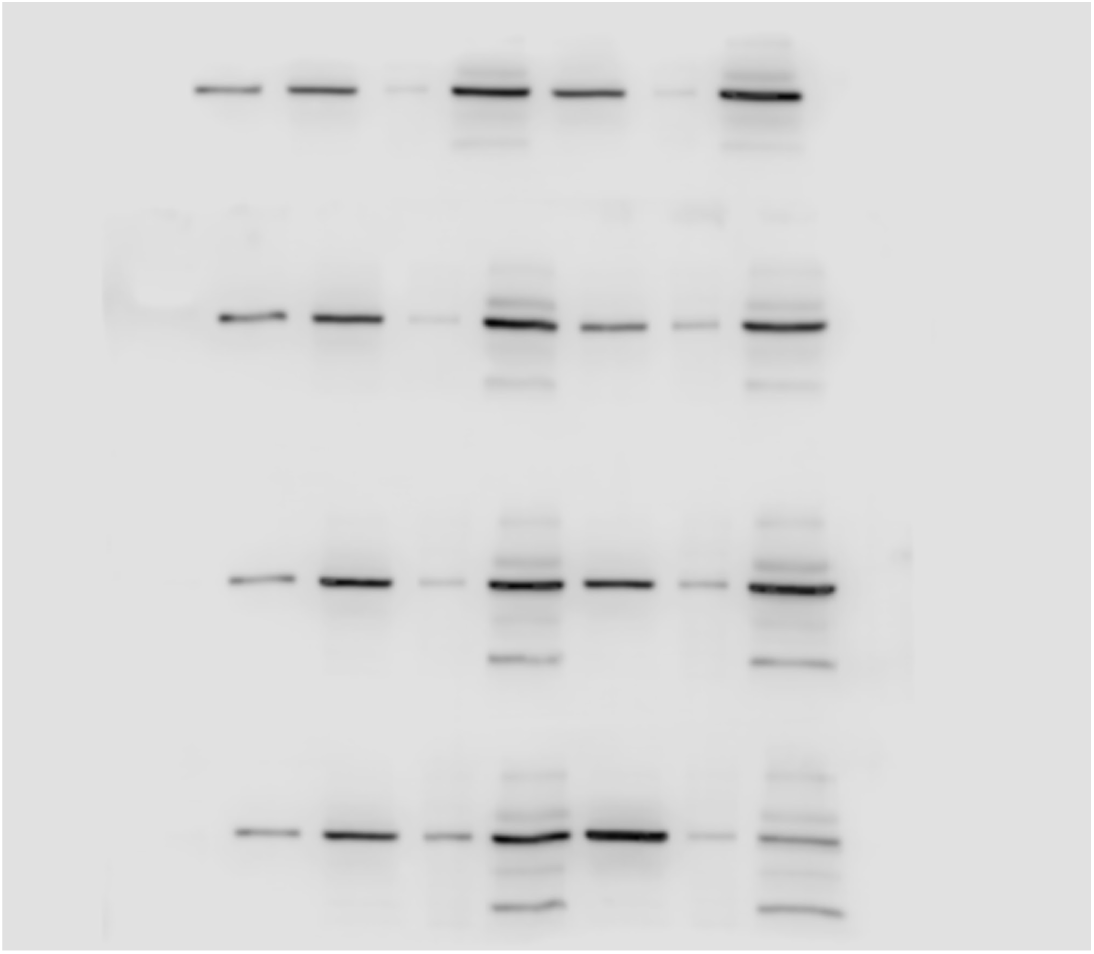
Taxol vs Control - Replicate 2. Western blot analysis of tubulin content present in the LSP, HSP, and HSS for Taxol-treated and control samples. The leftmost band of each row represents a 50 ng tubulin standard. Apart from that band, control samples are found in the left half of the 3rd row from the top, from left to right LSP, HSP, HSS. Taxol-treated samples are found in the right half of the 4th row from the top, from left to right: LSP, HSP, HSS.

**Figure S3:**
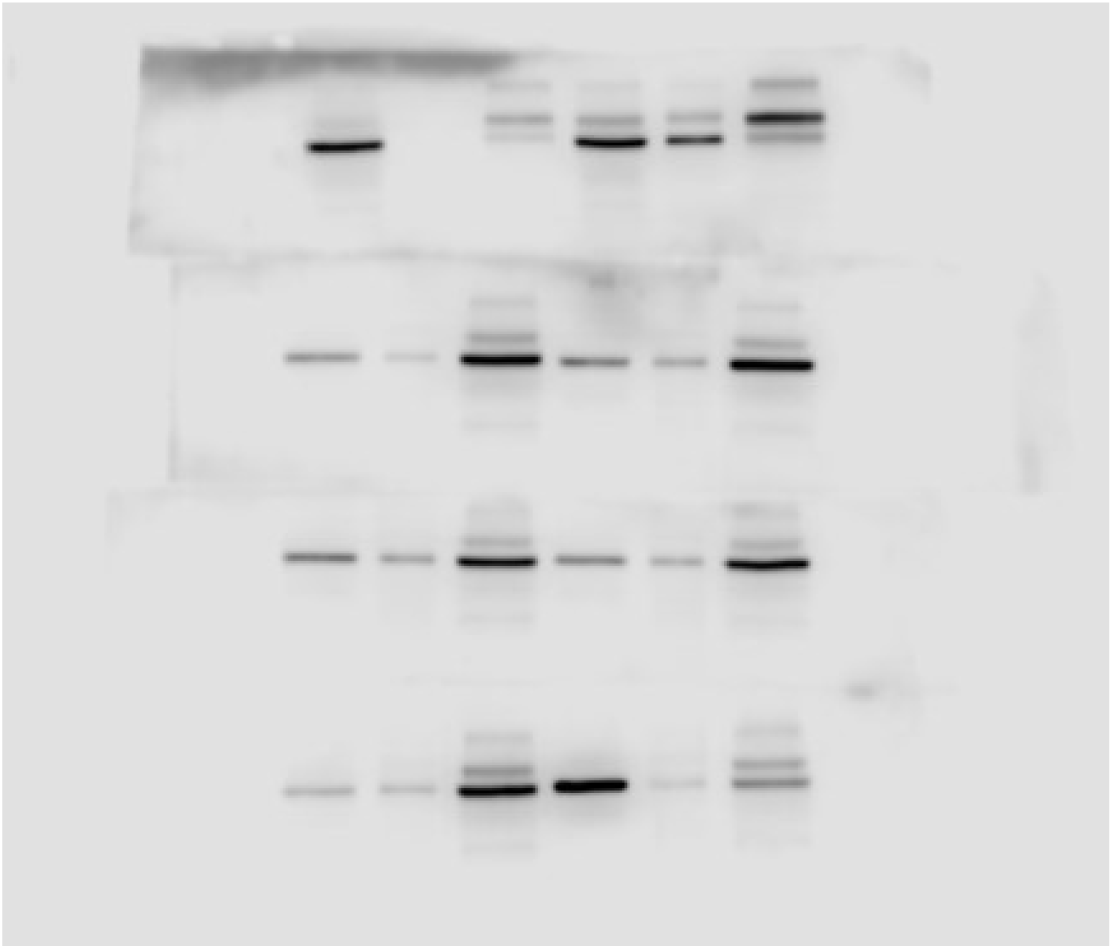
Taxol vs Control - Replicate 3. Western blot analysis of tubulin content present in the LSP, HSP, and HSS for Taxol-treated and control samples. Control samples are found in the left half of the 3rd row from the top, from left to right LSP, HSP, HSS. Taxol-treated samples are found in the right half of the 4th row from the top, from left to right: LSP, HSP, HSS.

**Figure S4:**
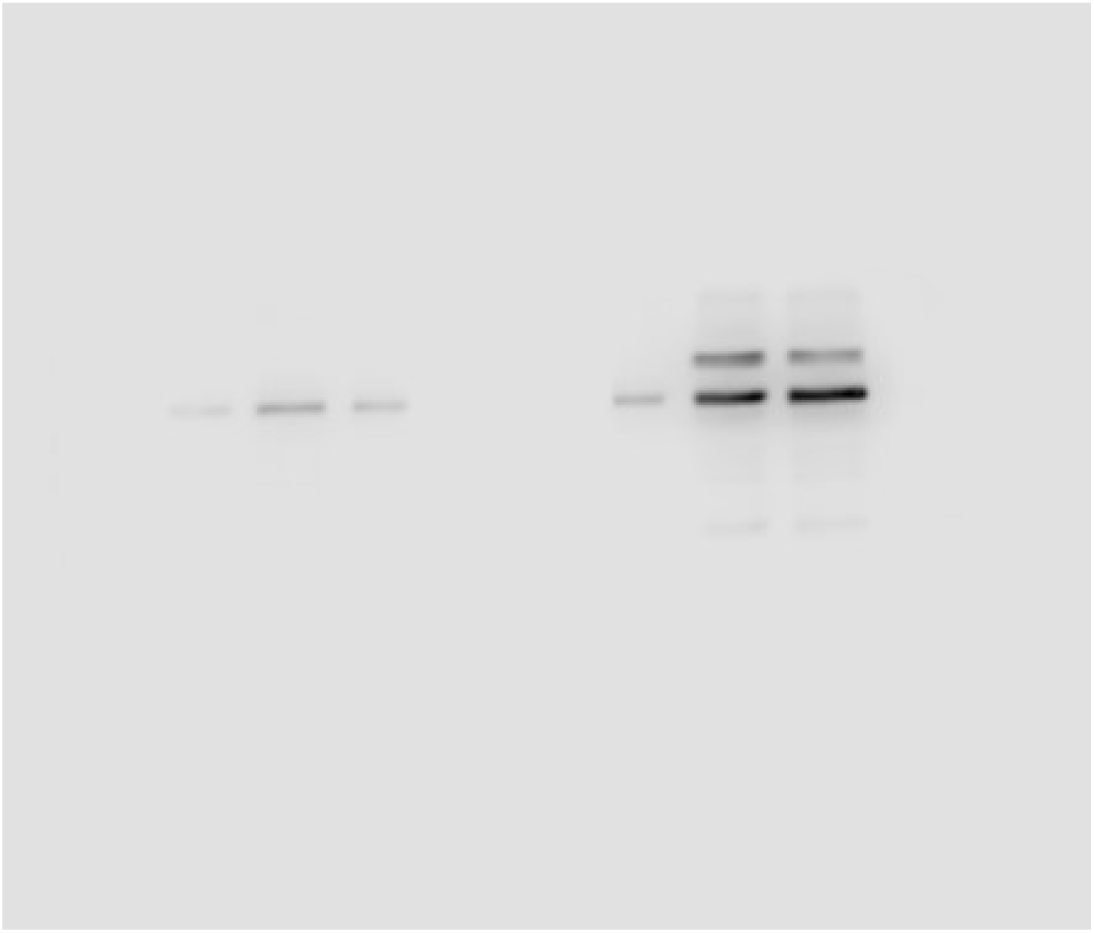
LIV vs Control - Replicate 1. Western blot analysis of tubulin content in LSP, HSP, and HSS for LIV-treated and control samples. From left to right, the bands are: control LSP, LIV LSP, control HSP, LIV HSP, control HSS, LIV HSS.

**Figure S5:**
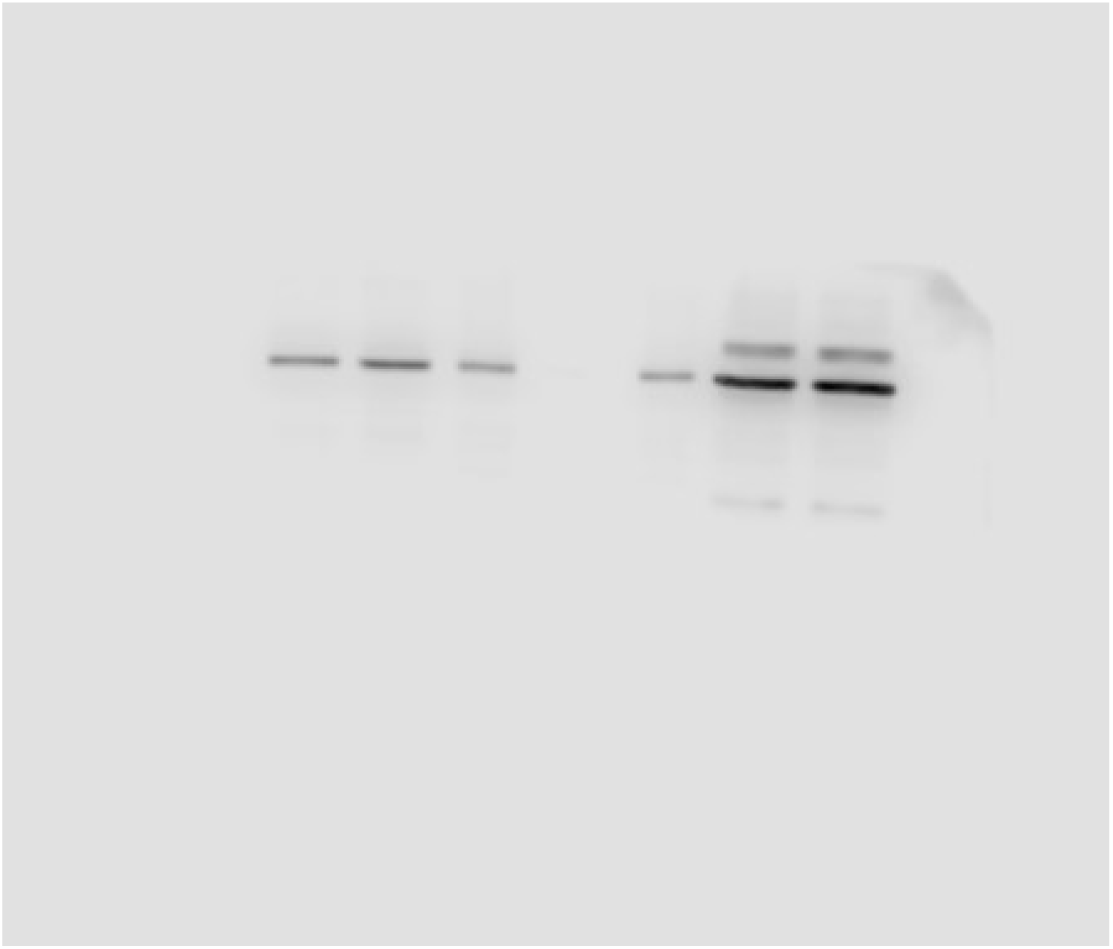
LIV vs Control - Replicate 2. Western blot analysis of tubulin content in LSP, HSP, and HSS for LIV-treated and control samples. From left to right, the bands are: control LSP, LIV LSP, control HSP, LIV HSP, control HSS, LIV HSS.

**Figure S6:**
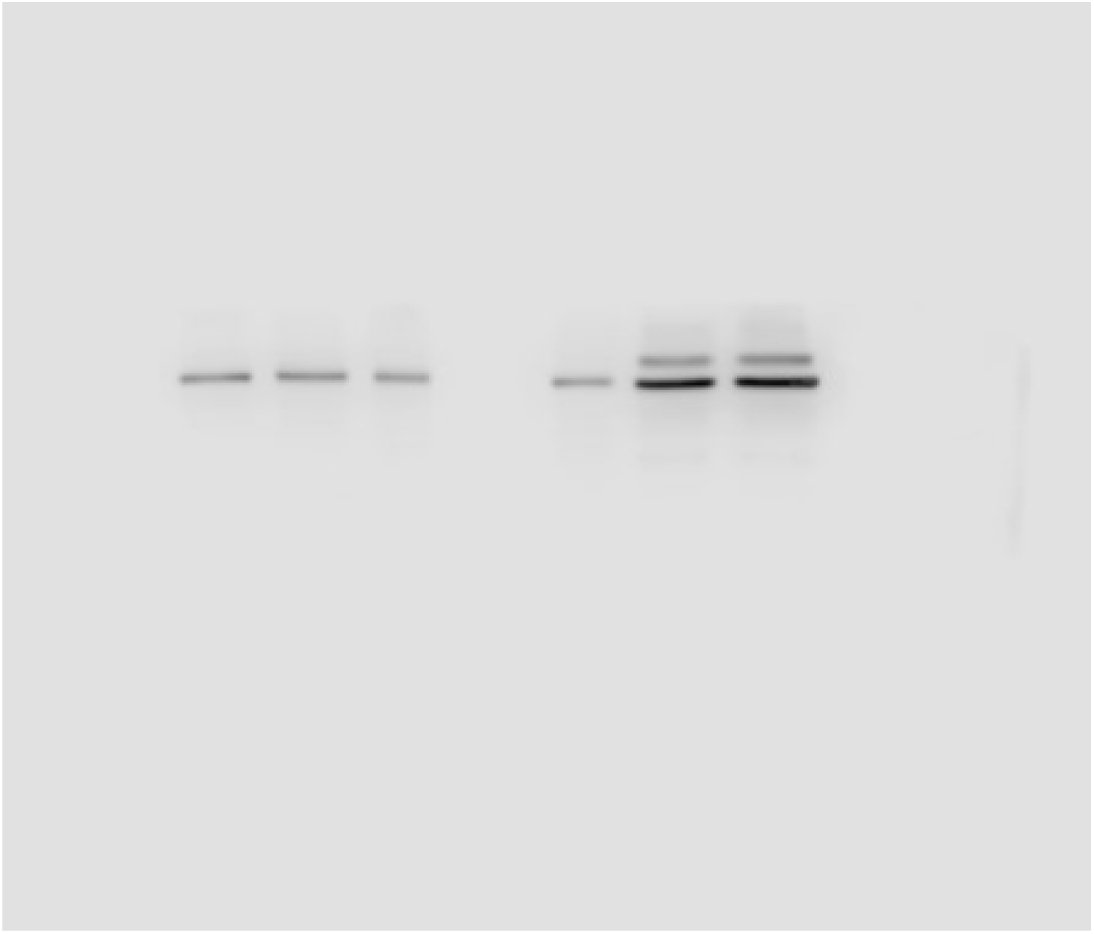
LIV vs Control - Replicate 3. Western blot analysis of tubulin content in LSP, HSP, and HSS for LIV-treated and control samples. From left to right, the bands are: control LSP, LIV LSP, control HSP, LIV HSP, control HSS, LIV HSS.

## References

1. Pittenger, M. F., et al. Mesenchymal stem cell perspective: cell biology to clinical progress. npj Regenerative Medicine. 4(1), 22; doi:10.1038/s41536-019-0083-6 (2019).

2. Uzer G., et al. Cell Mechanosensitivity to Extremely Low-Magnitude Signals Is Enabled by a LINCed Nucleus. Stem Cells. 33(6), 2063–76; doi:10.1002/stem.2004 (2015).

3. Rosenberg, N. The role of the cytoskeleton in mechanotransduction in human osteoblastlike cells. Human & Experimental Toxicology. 22(5), 271–4; doi:10.1191/0960327103ht362oa (2003).

4. Uzer, G., et al. Sun-mediated mechanical LINC between nucleus and cytoskeleton regulates betacatenin nuclear access. Journal of biomechanics 74, 32–40; doi:10.1016/j.jbiomech.2018.04.013 (2018).

5. Thompson, M., Woods, K., Newberg, J., Oxford, J. T., Uzer, G. Low-intensity vibration restores nuclear YAP levels and acute YAP nuclear shuttling in mesenchymal stem cells subjected to simulated microgravity. NPJ microgravity 6, 35; doi:10.1038/s41526-020-00125-5 (2020).

6. Touchstone, H., et al. Recovery of stem cell proliferation by low intensity vibration under simulated microgravity requires intact LINC complex *npj*. Microgravity 5, doi:10.1038/s41526-019-0072-5 (2019).

7. Uzer, G., Pongkitwitoon, S., Ete Chan, M. & Judex, S. Vibration induced osteogenic commitment of mesenchymal stem cells is enhanced by cytoskeletal remodeling but not fluid shear. Journal of biomechanics 46, 2296–2302; doi:10.1016/j.jbiomech.2013.06.008 (2013).

8. Pongkitwitoon, S., Uzer, G., Rubin, J. & Judex, S. Cytoskeletal Configuration Modulates Mechanically Induced Changes in Mesenchymal Stem Cell Osteogenesis, Morphology, and Stiffness. Scientific reports 6, 34791; doi:10.1038/srep34791 (2016).

9. Newberg, J., et al. Isolated Nuclei Stiffen in Response to Low Intensity Vibration. Journal of biomechanics, 111, 110012; doi:10.1016/j.jbiomech.2020.110012 (2020).

10. Akhmanova, A., Steinmetz, M. O. Control of microtubule organization and dynamics: two ends in the limelight. Nat Rev Mol Cell Biol. 16, 711–726; doi:10.1038/nrm4084 (2015).

11. Akhmanova, A., Steinmetz, M. O. Tracking the ends: a dynamic protein network controls the fate of microtubule tips. Nat Rev Mol Cell Biol. 9, 309–322; doi:10.1038/nrm2369 (2008).

12. Dogterom, M., Koenderink, G. H. Actin–microtubule crosstalk in cell biology. Nature Reviews Molecular Cell Biology. 20(1), 38–54; doi:10.1038/s41580-018-0067-1 (2019).

13. Gierke, S., Wittmann, T. EB1-Recruited Microtubule +TIP Complexes Coordinate Protrusion Dynamics during 3D Epithelial Remodeling. Current Biology. 22(9), 753–62; doi:10.1016/j.cub.2012.02.069 (2012).

14. Putnam, A. J., Schultz, K., Mooney, D. J. Control of microtubule assembly by extracellular matrix and externally applied strain. American Journal of Physiology-Cell Physiology. 280(3), C556–64; doi:10.1152/ajpcell.2001.280.3.C556 (2001).

15. Peister, A., et al. Adult stem cells from bone marrow (MSCs) isolated from different strains of inbred mice vary in surface epitopes, rates of proliferation, and differentiation potential. Blood 103, 1662–1668; doi:10.1182/blood-2003-09-3070 (2004).

16. Shord, S. S., Camp, J. R., Young, L. A. F. Paclitaxel decreases the accumulation of gemcitabine and its metabolites in human leukemia cells and primary cell cultures. Anticancer research. 25(6B), 4165–71; PMID: 16309212 (2005).

17. Applegate, K. T., et al. plusTipTracker: Quantitative image analysis software for the measurement of microtubule dynamics. Journal of Structural Biology. 176(2), 168–84; doi:10.1016/j.jsb.2011.07.009 (2011).

18. Ronneberger, O., Fischer, P., Brox, T. U-Net: Convolutional Networks for Biomedical Image Segmentation. Lecture Notes In Computer Science. 9351, 234–41; doi:10.1007/978-3-319-24574-4_28 (2015).

19. Kubitschke, H., et al. Actin and microtubule networks contribute differently to cell response for small and large strains. New Journal of Physics. 19(9), 093003; doi:10.1088/1367-2630/aa7658 (2017).

20. Hamant, O., et al. Developmental Patterning by Mechanical Signals in Arabidopsis. Science. 322(5908), 1650–1655; doi:10.1126/science.1165594 (2008).

